# FuzzyPPI: Human Proteome at Fuzzy Semantic Space

**DOI:** 10.1101/2023.05.24.541959

**Authors:** Anup Kumar Halder, Soumyendu Sekhar Bandyopadhyay, Witold Jedrzejewski, Subhadip Basu, Jacek Sroka

**Author notes:** Both shared 1^st^ authorthip.

## Abstract

Large scale protein-protein interaction (PPI) network of an organism provides key insights into its cellular and molecular functionalities, signaling pathways and underlying disease mechanisms. For any organism the total number of unexplored protein interactions significantly outnumbers all known positive and negative interactions. For Human, all known PPI datasets, contain only ∼ 5.61 million positive and ∼ 0.76 million negative interactions, that together is ∼ 3.1% of potential interactions. Moreover, conventional PPI prediction methods produce binary results. At the same time, recent studies show that protein binding affinities may prove to be effective in detecting protein complexes, disease association analysis, signaling network reconstruction, etc. Keeping these in mind, we present a fuzzy semantic scoring function using the Gene Ontology (GO) graphs to assess the binding affinity between any two proteins at an organism level. We have implemented a distributed algorithm in Apache Spark that computes this function and used it to process a Human PPI network of ∼ 180 million potential interactions resulting from 18 994 reviewed proteins for which GO annotations are available. The quality of the computed scores has been validated with respect to the available *state-of-the-art* methods on benchmark data sets. The resulting scores are published with a web-server for non-commercial use at: http://fuzzyppi.mimuw.edu.pl/.

## 1 Introduction

Proteins are involved in various biological functions in the cell through interactions with other proteins. Such interactions are often modeled as graphs with proteins as nodes and interactions as binary edges. These graphs are widely called protein-protein interaction (PPI) networks [1],[2].

A large scale PPI network of an organism provides valuable clues for understanding cellular and molecular functionalities [3], reconstructing signaling pathways [4], [5], multi-molecular complex detection [2], disease associations analyses [6], [7], drug discovery [8], [9], [10], We assume that the annotation ofetc. Tremendous effort has been invested into developing in-vitro experimental methods to extract positive PPIs [11], [12], [13], [14]. However, these experimental methods are able to produce only fraction of the PPI network of an organism while being costly and time-consuming.

Several computational approaches have been introduced to support PPI prediction [15], [16], [17], [18], [19], [20], [21], [22], [23]. The effectiveness of many of the *in silico* PPI prediction methods depends heavily on the selection of the positive and negative datasets during the training process. While biologically validated positive PPI datasets are available, negative databases are scarce [24], [25]. Therefore, *in silico* curation of negative datasets is an interesting research problem for the bioinformatics community.

In addition, if we consider an interactome with respect to any given organism, the total number of unexplored protein interactions significantly outnumbers the known positive/negative interactions, even if the ones obtained from *in silico* methods are considered.

For example, in Human proteome dataset from UniProt database [26], 20 350 reviewed human proteins result in ∼ 207 million potential interaction possibilities. However, so far we have information about ∼ 5.61 million positive and ∼ 0.76 million negative interactions, including both computational and experimental data. Together this is ∼ 3.1% of the interaction possibilities in the Human interactome. Therefore, there is a large number of unexplored interactions that needs to be assessed. One of the major challenges here is the cost of experimental PPI detection in the *in vitro* case and the sheer size of the complete interactome and lack of supporting biological data in the *in silico* case.

Finally, conventional PPI prediction methods generate binary interactions, whereas recent studies show that weighted interactions are proffered in pathway and/or disease analyses [27]. As has been shown in [28] protein-protein binding affinities may be effectively assessed with the help of the associated ontological networks.

### 1.1 Gene Ontology based model

In this paper we asses the probability of interaction between proteins based on the knowledge from Gene Ontology (GO) [29] knowledgebase, which is the world’s largest source of information on the functions of genes. GO is controlled and structured vocabulary of ontological terms that describe information about protein’s localization within cellular component (CC), participation in biological processes (BP) and association in molecular function (MF). GO terms are grouped into three independent direct acyclic graphs (DAGs) where nodes represent specific GO terms and the links among nodes represent different hierarchical relationships (see Figure 1). The relationships used in this experiment are: ‘*is_a*’, ‘*part_of* ‘, and ‘*has_part*’. A schematic diagram of GO-graph is shown in Figure 1.

**Fig. 1.**
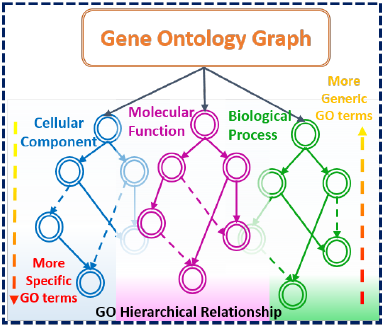
Schematic diagram of Gene Ontology details. Ontologies are organised in 3 hierarchical relationship graphs.

Each of the three subgraphs has a single most generic GO term, that can be considered the root in the sense that all the other nodes of that subgraph are reachable from it. The root nodes are GO:0005575, GO:0003674 and GO:0008150 for CC, MF and BP, respectively. Because of that, tree oriented notions are often generalized and used with GO subgraphs. For example a specific GO term is considered a descendant of the more generic GO terms from which it has incoming edges [30], [31], [32].

The UniProt-Gene Ontology Annotation (UniProt-GOA) database (see http://geneontology.org/docs/go-consortium/) includes information about GO annotation of proteins in the UniProtKB dataset. For any protein, GO annotations are broadly classified into two groups based on inference evidence: inferred from electronic annotation (IEA) and the manually reviewed (non-IEA). The non-IEA evidences includes data inferred from Experiment (EXP), Direct Assay (IDA), Expression Pattern (IEP), High Throughput Direct Assay (HDA), etc. (for details see http://geneontology.org/docs/guide-go-evidence-codes/). The inclusion of IEA which provides ∼ 25% of all annotations enhances coverage. In practice most of the semantic assessments with GO are determined in two orientations, one being ‘with-IEA’ and the other ‘without-IEA’. This results in a total of six semantic assessment scores for each pair of proteins, *i.e*., IEA and non-IEA scores for CC, BP and MF, respectively. In this paper we compute such scores and combine them into a single semantics assessment score, ranging from 0 to 1, to which we refer to as *fuzzy semantic score of binding affinity*.

At the time of this writing, the GO dataset contains 44 579 annotations (release-version:*02-2020*) of which 29 267 are BP, 11 126 MF and 4 186 CC. The GO terms annotated to proteins can be used to infer functional relationship between them. To compute the binding affinity between two proteins we need to estimate the semantic similarity scores of all the GO term pairs, where the first term is annotated to one of the proteins and the second to the other. Semantic similarity computation for any GO pair by exploring three different GO relationship graphs is a time consuming task. Thus there is a strong need for a method that can handle large data and is easy to distribute.

In this paper, we propose an efficient algorithm to compute the fuzzy binding affinity for large scale PPIs. We then implement it using a high throughput parallel architecture based on Apache Spark. Spark is a popular platform for large scale data analysis [33], [34]. Its main advantages over previous frameworks like Hadoop are: expressive and high level API, and the ability to keep intermediate results in memory between computation phases. The latter saves disk I/O and results in a huge efficiency gain. Furthermore, Spark has sub-modules for data analytics, graph processing, machine learning and streaming so combining such applications in one project does not introduce any integration overhead and at the same time Spark core optimizations, of which there are many, are applied on the end to end data processing pipeline.

### 1.2 Related Work

PPI prediction is an established and active domain. Here we provide an overview of recent developments and position our work in their context.

Several methods have been developed to measure the semantic similarities between protein/gene pairs. The existing approaches can be classified into three broad categories based on the information that has been used from the GO relationship graphs. These are node-based, link or edge-based, and hybrid methods.

Edge-based methods rely on the distance between two GO terms in the GO graph [35], [36], [37]. Usually, the distance is computed as the number of edges in the shortest graph path between two terms or as an average over all paths. Such distance can easily be normalized and converted into a similarity measure. Another way to compute similarity is to check the depth of the first common ancestor of the nodes. The deeper from the root of the subgraph the common ancestor is, the more the nodes have in common. Unfortunately, the nodes and edges in the GO subgraphs are not distributed uniformly, nor edges at the same level in the ontology correspond to the same semantic distance between terms [38]. In [39] such issues were addressed by weighting edges differently according to their hierarchical depth, or taking into account node density and link type. Yet, this does not fully solved the problem and edge-based strategies and edge-based approaches are considered ineffective in terms of semantic assessment [28].

Node-based approaches utilize the properties of the pair of GO terms themselves and their ancestor or descendant nodes [40], [41], [42], [43]. Sometimes the concept of information content (IC) is used [44], which is a measure of how specific and informative a GO term is. Resnik [40] has defined the semantic similarity as the IC of the most informative common ancestor (MICA) of two GO terms. The MICA and the lowest common ancestor node refer to the same ancestor of two GO terms where the MICA is presented in the context of searching common path between GO terms, and the latter is presented in the context of IC of GO terms. In semantic similarity computation, several methods have used the IC values of query proteins [41], [42]. Another IC based approach has been proposed by Schlicker *et al*. [43] where the relevance similarity measure has been defined using the location of the query GO terms in the DAG by considering the properties of MICA [45]. Mazandu and Mulder [46] have proposed a method that normalizes the IC-based semantic similarity to 1 when measuring the similarity between the same GO terms. A new approach, *GraSM* has been introduced in [47] to avoid the over-reliance on MICA. It is designed in such a way that it can be applied to any IC-based methods where the semantic similarity is calculated by the average IC of the disjunctive common ancestors (DCAs). The DCAs are identified by the number of distinct paths from the query GO terms to MICA [47].

The IC-based methods have an obvious advantage, as they use IC to indicate the specificity of a GO term and are because of that free from the problems of nonuniform semantic distance and edge density. However, corpora-dependent IC calculation can cause problems. In general, IC-based approaches suffer from two major issues. First, the IC calculation depends on the annotation corpora set, as same GO term may have different IC values when different corpora are used [48]. Second, the IC is biased by the research trend [38], as the GO terms related to popular fields tend to be annotated more frequently than the ones related to less popular fields and the annotation of some terms may still be missing in the corpus [45]. These issues largely degrade the overall performance and effectiveness of methods that only use ICs.

To overcome the limitations of the IC-based approaches, many hybrid methods have been developed that consider both edges and nodes in the DAG. Wang *et al*. [20] have proposed a hybrid GO-universal method that calculates the semantic similarities based on the topology of GO DAG. It takes into account the topological position characteristics in the GO subgraphs and considers the number of children terms instead of the frequency of terms from the annotation corpus. GO-universal defines the topological position characteristic of the root to be 1 and calculates the topological position characteristic of a non-root GO term by using a ratio based on the number of children of all ancestor GO terms [49]. A hybrid structural similarity based method has been proposed by Nagar and Al-Mubaid [50] using the shortest path plus either IC generated from corpora or structure-based IC generated from DAG. In a recent work, Dutta *et al*. [51] have proposed a hybrid semantic similarity measure between two GO terms based on a combination of topological properties of the GO graph and average IC of the DCAs of the GO terms.

### 1.3 Contributions

Although the GO based methods are popular in assessing PPI binding affinity, one of the major challenges lies in managing the computational overhead. For example, in Human PPI network the total number of protein interactions to be explored is in the order of hundreds of millions. This has been the primary concern for most of the prior studies [45], [47], [49], [51], which concentrated on a smaller subset of the complete dataset. Due to our distributed architecture and optimized algorithm we have eliminated this problem. We use the complete human proteome and the underlying GO annotations for efficient assessment of the PPI binding affinity and propose a novel fuzzy semantic score. The outcome of our approach is not limited to quantitative analysis. The fuzzy semantic network helps to understand different biological mechanisms such as in protein complex module identification and functional analysis, high quality data identification for PPI model, PPI interaction analysis, etc.

Our contributions in this paper are summarized as follows:

- Design of a fuzzy semantic score of binding affinity using the GO network to assess the binding affinity between any two proteins at an organism (Homo sapiens in our work) level.
- Implementation of the underlying algorithm using a Spark based parallel architecture and using it to process a Human PPI network of ∼ 180 million potential interactions resulting from 18 994 reviewed proteins for which GO annotations are available.
- Construction of fuzzy semantic network at proteome level from the above designed binding affinity function and extraction of meaningful biological insights.
- Validation of the developed method with respect to the available *state-of-the-art* methods on benchmark datasets.
- Development of a FuzzyPPI web-server with pre-computed PPI affinity scores, available freely for non-commercial use, at: http://fuzzyppi.mimuw.edu.pl/.

The rest of the paper is organized as follows. In Section 2, we discuss the datasets used for the experiment and also for the benchmark evaluation. We present the overall methodology in Section 3 and the design of the fuzzy semantic score of binding affinity in section in Section 3.3. Then, in Section 3.5, we explain the design of the parallel architecture and the underlying algorithms. Experimental results are discussed in Section 4, followed by the conclusion in Section 5.

## 2 Dataset

In this section we describe how the input data for our experiment was prepared.

We have used two datasets. The first one, obtained from UniProt, contained information about human proteins and their GO annotation. We have used it as a base for our predictions. The second one, has been obtained by combining several popular datasets and composed of experimentally validated information about positive and negative interactions between human proteins. We have used it as a benchmark for our analysis.

The details of these dataset are discussed in Subsections 2.1 and 2.2, respectively.

### 2.1 Proteome data

The UniProt knowledgebase (UniProtKB) [26] is the combined and uniform collection of proteins from Swiss-Prot [52], TrEMBL [53], and PIR-PSD [54]. This database maintains a bi-directional cross references between *“PIR-PSD”* and *“Swiss-Prot + TrEMBL”* protein entries. The main objective of this database is to maintain a single entry and to merge all reports for any particular protein [52]. Thus, it reflects a complete structure of protein repository for different organisms where many of the entries are derived from genome sequencing projects.

As UniProt is highly popular, with over 0.4 million unique visitors of its web site per month [55] and over 20 000 total citations of its main publications, a thorough analysis and annotation by biologists ensures the quality of this database.

In UniProt, human proteome (*id:UP000005640*) contains 20 350 manually annotated human proteins. Among these, 18 994 proteins are annotated with GO annotations which is ∼ 93% of reviewed proteins. In our experiment, the fuzzy binding affinity is assessed for the resulting 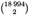 ∼ 180 million pairs of unique proteins.

UniProt allows to export protein data together with annotations in a text format presented in first part of Figure 2. For each protein with UniProt id (e.g. P29274) and annotated to it GO term (e.g. GO:0030673) there is information about GO type (C, F or P) and evidence type (IEA, non-IEA). We encode this in a hierarchical format as shown in Figure 2 and store as a Parquet file which is a binary and columnar format integrated with Spark with optimizations that allow significant performance improvement.

**Fig. 2.**
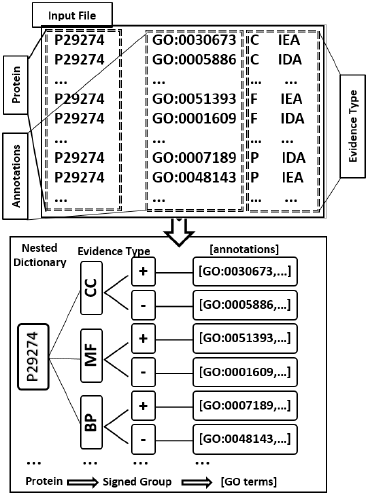
Transformation of UniProt text format to hierarchical representation of Protein-GO annotations

### 2.2 Protein interaction data for benchmarking

To prepare the benchmark dataset we have used experimentally validated human protein interactions from several popular datasets such as HIPPIE [56], STRING [57], BioGRID [58], DIP [59], HuRI [60] for positive data and Negatome 2.0 [24], Trabuco *et al*. [25] for negative data. The statistics of the interaction data are presented in Table 1.

**TABLE 1.** Database wise proteome level statistics of interactions of UniProt reviewed proteins. (Green and yellow colors represents positive and negative interaction datasets, respectively.)

| Database | Existing Intr. |  | Unexplored Intr. |  |
| --- | --- | --- | --- | --- |
|  | Individual | Total | Total | (in %) |
| DIP [59] | 4 652 | 5 611 787 | 200 680 877 | 96.9% |
| HuRI [60] | 64 711 |  |  |  |
| BioGRID [58] | 104 509 |  |  |  |
| HIPPIE [56] | 342 996 |  |  |  |
| STRING [57] | 5 415 071 |  |  |  |
| Negatome 2.0 [24] | 1 482 | 758 411 |  |  |
| Trabuco *et al.* [25] | 756 994 |  |  |  |

We have combined them into a single dataset with ternary information (interact, do not interact, unknown). We consider that there is an interaction between two proteins if there was evidence for that in any of the positive datasets and no evidence for the lack of interaction in all of the negative datasets. The other way around, we consider that there is no interaction between two proteins if there was evidence for that in any of the negative datasets and no evidence for interaction in all of the positive datasets. We also distinguish gold sample of interactions for which we have high confidence. The resulting combined dataset has information about 5 107 321 positive interactions, 730 122 negative interactions and 18 295 out of the total 18 994 proteins are somehow accounted. For the gold sample the statistics are: 361 076 positive interactions, 182 667 negative interactions and 15 261 proteins. The statistics of interactions for individual proteins are presented in Figure 3.

**Fig. 3.**
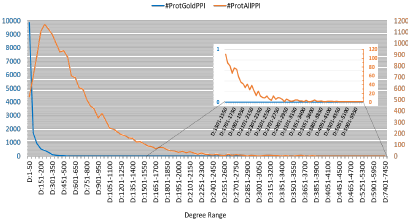
Degree-range wise plot of positive interactions.

The databases have different scoring schemes to quantify the interaction quality. For example for BioGRID the values are in range [− ∞, + ∞], while for HIPPIE in [0,1]. This is summarized in Table 2 where we also specify the thresholds for considering two proteins to interact and for including the pair in the gold sample. For other databases a binary information is present but the data is grouped in high and low confidence groups. For example, for DIP there are two groups core and non-core and we consider both of them as the source of evidence for interactions, but only core as the source of evidence for high quality interactions. Sometimes, as is the case for HuRI, one group is a superset of the other, high quality group. Finally, GoldPos and AllPos interaction datasets are constructed by considering the union of individual gold data and all data respectively from the above described five positive interaction databases. The degree-range specific density plot of all proteome is shown in Figure 3. Recall, that for negative interactions we also require that no positive interaction exists in any of the datasets. Furthermore, for gold standard negative interactions we also forbid in silico positive interactions. In Negatome *2.0*, [24] the negative data are categorised as gold based on the selection strategy as of Manual stringent (MS). For the dataset by Trabuco *et al*. [25], high quality negative set is selected by removing the positive interactions that have any type (*in-vivo, in-vitro* and *in-silico*) of evidence of positive interaction (AllPosR) while all negative set is constructed by removing the interactions that belong to the gold quality positive (GoldPos) interactions. Note that the AllPosR set includes in-silico interactions where AllPos excludes in-silico.

**TABLE 2.** Details of benchmark interaction data selection (Green and yellow colors represent positive and negative interactions datasets, respectively.

| Database | Score range/groups | All-Interactions filter | Gold Quality filter |
| --- | --- | --- | --- |
| BioGRID [58] | $[-\infty, +\infty]$ | $\geq 0$ | $\geq 10$ |
| HIPPIE [56] | $[0, 1]$ | $\geq 0.75$ | $\geq 0.9$ |
| STRING [57] | $[0, 1000]$ | $\geq 150$ | $\geq 900$ |
| DIP [59] | *core(C)*,<br>*non-core(NC)* | C+NC | C |
| HuRI [60] | *HuRI(H)*,<br>*HuRI-Union(HU)* | HU | H |
| Negatome 2.0 [24] | *Manual(M)*,<br>*Manual-stringent(MS)* | M+MS | MS |
| Trabuco *et al.* [25] | Negative ($N^T$) | $N^T - \text{GoldPos}$ | $N^T - \text{AllPosR}$ |

## 3. Methods

Building on the model from our previous work [51] in this section we formally define how the fuzzy semantic network can be used to assess the interaction affinity of any two proteins with GO annotations. The semantic similarity (SS) between any two proteins is estimated based on the similarities of pairs of GO terms such that one GO terms annotates first protein and the other the second. The pairs are considered independently for each of the GO subgraphs, *i.e*., we consider only pairs of terms from the same GO subgraph. For each GO subgraph, semantic similarity is computed in two ways based on the evidence type of annotations, *i.e*., for IEA and nonIEA which are denoted as + and -, respectively. A Spark-based parallel implementation of this scheme has been designed by leveraging its high throughput architecture for large scale proteome-level interaction analysis. The implementation details of the fuzzy function and parallel processing are presented in the following section. Here we define what is being computed.

### 3.1 GO subgraphs

We will use the standard notation to denote a directed graph *G* = ⟨*V, E*⟩, where the first element of the tuple *V* is the set of nodes and the second *E* the set of edges between the nodes, i.e., *E* ∈ *V* × *V*.

Let *CC* = ⟨*V*_*CC*_, *E*_*CC*_⟩, *MF* = ⟨*V*_*MF*_, *E*_*MF*_⟩ and *BP* = ⟨*V*_*BP*_, *E*_*BP*_⟩ be the GO subgraphs for cellular component (CC), molecular function (MF) and biological process (BP), respectively. By assumption *V*_*CC*_ ∩ *V*_*MF*_ = *V*_*CC*_ ∩ *V*_*BP*_ = *V*_*MF*_ ∩ *V*_*BP*_ = ∅ from which it follows that *E*_*CC*_ ∩ *E*_*MF*_ = *E*_*CC*_ ∩ *E*_*BP*_ = *E*_*MF*_ ∩ *E*_*BP*_ = ∅.

We also assume that the *anc*-*or*-*self* and *desc*-*or*-*self* functions are defined for each graph node with the standard meaning.

### 3.2 GO annotation

We assume that the annotation of proteins with GO terms (see Figure 2) is available as, such that for each protein *p*, a GO subgraph *SG* ∈ {*CC, MF, BP*}, and the evidence type annotation *iea* ∈ {+ −}, (IEA or nonIEA, respectively) it returns the set of all GO terms from *V*_*SG*_ that are annotated with specified IEA/nonIEA annotation type *iea*. For example, as shown in Figure 2, 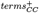 (*P* 29274) = *GO* : 0030673, … while 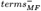 (*P* 29274) = *GO* : { 0001609}, Similarly we define *prot*^*iea*^(*t*), such that for each protein *t* and evidence type annotation *iea* ∈ {+ −}, (IEA or nonIEA, respectively) it returns the set of all proteins that are annotated with term *t, i.e* 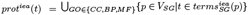.

### 3.3 Fuzzy Semantic Scoring for proteins

The main intuition that we are going to follow is that two proteins having the same functions are likely to interact to complete the function. Also the more of functions they share and the more specific the common functions are the higher the interaction chance is. The functions of proteins are concluded from GO annotations that these proteins have. Different GO subgraphs represent different types of functionality and are examined separately and then combined. This way numerous annotations from one GO subgraph do not dilute possibly less numerous but highly shared annotations in other subgraphs.

We first formally define the *semantic score* of binding affinity for pairs of proteins and then normalize it into *fuzzy semantic score*. The scoring of proteins is based on the scoring of the pairs of GO terms annotated to them that is defined in Subsection 3.4. For any pair of proteins *p*_*a*_ and *p*_*b*_ semantic score (SS) of binding affinity is defined as:

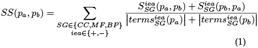

Where 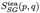 is the similarity of proteins *p* and *q* within the GO subgraph SG, based on the IEA/nonIEA annotation type *iea*, and is defined as:

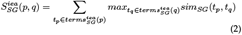

and *sim*_*SG*_(*t*_*p*_, *t*_*q*_) is the semantic similarity for a pair of GO terms that is defined in Subsection 3.4.

We normalized the semantic score with the max-min normalization. It linearly transforms SS into fuzzy semantic score (FSS):

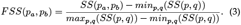

Note that FSS results are limited to [0, 1] and can be viewed as probability of interaction between proteins *p* and *q*.

### 3.4 Semantic Similarity for GO terms

In the following Subsections we define *MDMS*_*SG*_(*t*_*i*_, *t*_*j*_) to be the maximal difference in membership strengths (see Subsection 3.4.1) of the GO terms *t*_*i*_, *t*_*j*_ to natural clusters covering of subgraph SG and *SIC*(*t*_*i*_, *t*_*j*_) to be the shared information content (see Subsection 3.4.3) between the target GO-terms. With those we define the *semantic similarity* of a pair of GO terms that come from the same GO subgraph.

#### Definition 3.1

(semantic similarity). For a GO subgraph *SG* and *t*_1_, *t*_2_ ∈ *V*_*GO*_ the semantic similarity of *t*_1_ and *t*_2_ is defined as: *sim*_*SG*_(*t*_1_, *t*_2_) = (1 − *M D M S*_*SG*_(*t*_1_, *t*_2_)) * *SIC*(*t*_1_, *t*_2_).

Note that the semantic similarity of two terms is highest if at the same time their maximal difference in membership strengths is small and shared information content is high.

#### 3.4.1 Maximal difference in membership strengths

The maximal difference in membership strengths *MDMS*_*SG*_(*t*_*i*_, *t*_*j*_) is computed in respect to the set *CC*_*SG*_ of cluster centers of SG, which we define in the next subsection. For each cluster center we can compute how close other nodes are to it, *i.e*., how strong is their membership to this cluster. Large differences in membership mean that nodes are not similar. Following [51] we use Gaussian to convert distance to the cluster center into similarity and compute the maximum difference of such similarities (membership strengths) over all cluster centers:

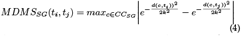

where *d*(*t*_1_, *t*_2_) is the smallest distances between nodes *t*_1_ and *t*_2_ taken in *V*_*SG*_ and in 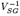. That is *d*(*t, t*) = 0 and for *t*_1_ ≠ *t*_2_ we define *d*(*t*_1_, *t*_2_) as the smallest *n* such that there exists a sequence of nodes *w*_1_, *w*_2_, …, *w*_*n*−1_ for which there exists a path (*t*_1_, *w*_1_), (*w*_1_, *w*_2_), …, (*w*_*n*−1_, *t*_2_) ∈ *V*_*SG*_ or a path 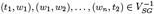.

#### 3.4.2 Natural cluster covering of a GO subgraph

In [51] the cluster centers *CC*_SG_ for each GO subgraph *SG* were chosen to be the nodes with high values of ratio of nodes reachable from them together with nodes from which they are reachable to the total number of nodes in the graph. We first briefly define the ratio and then explain how we improved upon the selection of cluster centers for this paper.

To formally define the ratio, we use transitive closure of the edge relation *E*^*^ defined as the set of all node pairs ⟨*u, v*⟩ ∈ *V* × *V* such that there exists a (*u, v*)-path in the graph, *i.e*., (*u, w*_1_), (*w*_1_, *w*_2_), …, (*w*_*n*_, *v*) ∈ *E*. We say that node *v* ireachable from *u* if ⟨*u, v*⟩ ∈ *E*^*^ and we denote the set of all nodes reachable from *u* as *E*^*^(*u*). With this the ration used to determine the cluster centers for any *v* ∈ *V* is defined as:

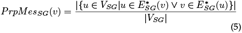

The ratio based method proposed in [51] and other works leads to selection of 49 cluster centers for CC, 32 for MF and 68 for BP. Here, we have tried centroid based clustering methods adapted to work on the respective graphs that consider the graph topology by minimizing the distance between cluster centers and cluster members. We have obtained the best results with the K-Medoids Clustering method [61]. We have started from the centers from [51] and we have added more centers using the k-medoids++ initialization method until each term was reachable from some center. This resulted in 149, 728 and 577 GO nodes as cluster centers from CC, MF and BP subgraphs respectively. The new sets of centers share 23 (CC), 19 (MF), and 28 (BP) common GO terms with Dutta *et al*. with 0.131 (CC), 0.126 (MF), and 0.045 (BP) Jaccard similarity. In gold standard positive and negative PPI dataset, our newly developed cluster centre selection algorithm has improved the overall performances (best AUC score: 0.780 at *μ* = 0.15) compared to Dutta *et al*. [51] (best AUC score: 0.764 at *μ* = 0.15) and in ALL-PPI dataset, the improvement is 1.5%.

#### 3.4.3 Shared Information Content (SIC)

Now we take into account how informative the common ancestors of a pair of terms are. The intuition is similar to the notion of term frequency–inverse document frequency (TFIDF) [62] where rare words tend to more correctly summarize a document than very common words. Similarly some terms that are commonly annotated to proteins are less useful in comparing them as opposed to rarely occurring terms. For that we define information content (IC) of a term in GO subgraph SG with respect to the evidence type annotation iea:

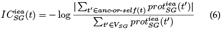

To obtain the SIC between a pair of GO terms, the average IC of all disjunctive common ancestors (DCAs) of the GO terms is computed [51]:

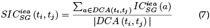

The DCA of any GO terms *t*_*i*_, *t*_*j*_ are those common ancestors *a* such that for a given difference of number of paths to *a* from *t*_*i*_, *t*_*j*_ they have the highest information content. Let *CA*(*t*_*i*_, *t*_*j*_) be the set of all common ancestors of terms *t*_*i*_ and *t*_*j*_ defined as *CA*(*t*_*i*_, *t*_*j*_) = *anc*-*or*-*self* (*t*_*i*_) ∩ *anc*-*or*-*self* (*t*_*j*_). Let *PD*(*a, t*_*i*_, *t*_*j*_) be the difference in number of (*a, t*_*i*_) and (*a, t*_*j*_) paths. The DCA is defined as follows:

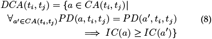

### 3.5 Parallel Implementation with Spark

In this section we present our distributed algorithm and its Spark implementation of fuzzy semantic scoring function. Given a set of proteins, GO subgraph *SG* and annotation type *iea* (either IEA or nonIEA) that defines GO terms assigned to the proteins, the algorithm computes 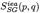 as defined in 3.3. The algorithm is divided into four phases that include: one phase of preprocessing of the input data, two phases of precomputation of values that otherwise would be computed many times and the final computation using the precomputed values.

#### 3.5.1 Preprocessing: Hierarchical representation of Protein-GO annotation data

In the first phase, the annotation data is parsed and preprocessed into hierarchical data structure representing *iea* annotations. The information about 18 994 proteins and their annotation with ∼ 266 million GO terms is stored as a text file (sample representation shown in Figure 2) and is considered as an input to the pre-processing step. The information is organized into a hierarchical structure (nested dictionary) in which each protein id is mapped to list of its GO terms grouped by GO subgraphs and annotation types. This structure is used to extract the dataset of proteins with their respective GO terms for selected *SG* and *iea*, which is needed in subsequent algorithm phases. As the size of the input data is small, this phase does not have to be distributed.

#### 3.5.2 PreComputation-I: Finding unique GO term pairs from all protein pairs

Our goal is to compute Fuzzy Semantic Score for each protein pair as defined in Section 3.3. For that we need to calculate and combine semantic similarities of all GO term pairs for the protein pair. As the number of protein pairs is quadratic in respect to the number of proteins, this computation is substantial and should be distributed. The pseudocode is presented in Algorithm 1. Naive implementation would repeat computation of semantic similarities for GO term pairs for different protein pairs. In order to avoid this, we pre-compute semantic similarities for all pairs that appear at least once for any protein pair. Using Spark, we distribute the set of proteins along with their GO terms on the cluster in the form of a Spark distributed collection DataFrame. Then, we leverage the Spark built-in mechanism to obtain the Cartesian product of the DataFrame with itself. We keep only those rows of the product in which the first protein is lexicographically smaller than the second one to get rid of duplicate combinations. Then, using Spark’s flatMap transformation, we obtain DataFrame of all GO term combinations for all proteins. Also here we make sure not to distinguish the pairs where the same elements are in different order. Finally, we remove duplicate pairs with Spark distinct transformation, which is implemented efficiently with grouping and combining on the map side of the shuffle. The resulting set of unique GO term pairs for human proteins is small enough (8 152 365k pairs before de-duplication and canonical ordering, 73 210k after de-duplication) to be collected to a single machine.

##### Algorithm 1 UniqueGOPairComputation

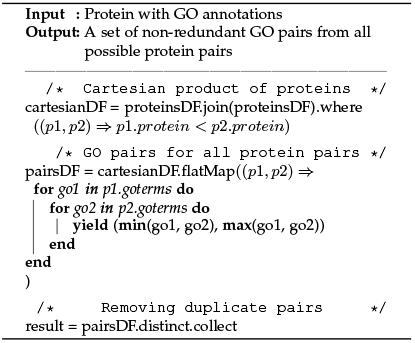

#### 3.5.3 PreComputation-II: Computation of the semantic similarity for unique GO term pairs

Next, we can compute the semantic similarity score, as defined in Subsection 3.4, for all unique GO term pairs obtained in the previous step. The computation for each pair is independent so it can be easily distributed if the number of GO term pairs grows large. Thanks to limiting of the number of pairs in the previous step, for human proteins we need to compute score for only 73 210k pairs which can be completed in half an hour by a sequential, centralized algorithm and with further linear speedup due to distributing. The resulting similarities take little more than 1GB and can be easily broadcast to every node in the cluster for the next step.

To compute semantic scores for all the GO term pairs, we first compute *SIC* scores (see 3.4.3) for each pair, then *MDMS* (see paragraph 3.4.1) for each pair, and finally we combine them to get the semantic similarity.

##### Algorithm 2 numberOfPaths

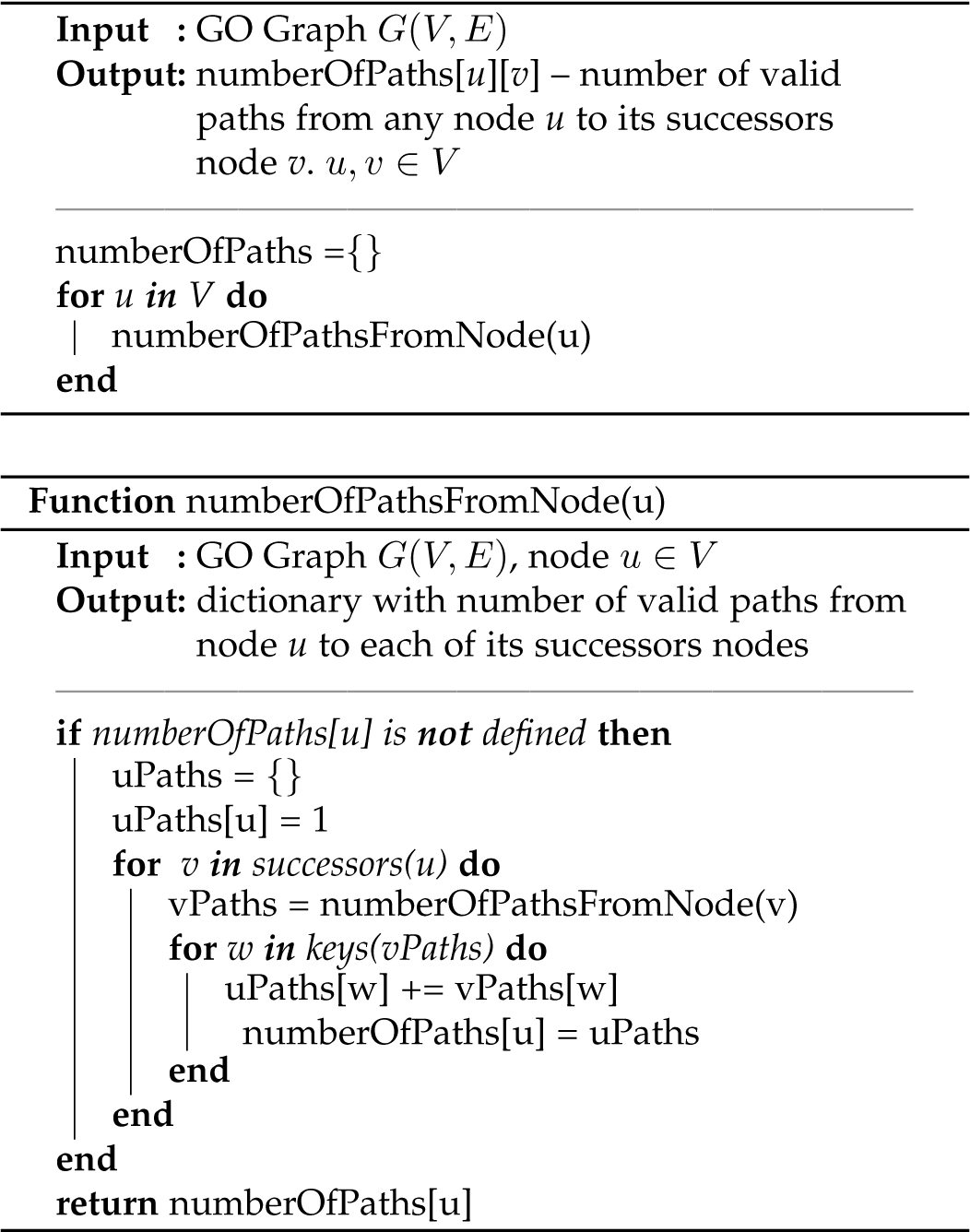

We start by explaining how we compute the SIC score for each GO term pair. Based on the GO subgraph *SG*, for each pair (*t*_1_, *t*_2_) of GO terms present in the subgraph we precompute the number of different paths from *t*_1_ to *t*_2_. This can be done in time quadratic to the graph size using Depth-first Search. The result can be stored in a sparse form as a map that for each node stores a map from another node into the number of paths between them. The lack of entry in the map for a given node pair denotes that there are no paths between the pair. The relevant pseudocode is presented in Algorithm 2.

We also precompute the information content 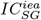 (see paragraph 3.4.3) for all the GO terms. For that, once we compute all the paths for each GO term as well as *IC* scores, we can compute *SIC* score of each GO term pair. We use the pre-computed structure with the numbers of paths to iterate over all of the descendants of a term, along with the number of paths to it. The iteration is performed over all the combinations of two descendants excluding the pairs that are not present in unique GO term pairs set refereed in paragraph 3.5.2 for reducing the redundancy. The relevant maximum 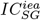 value is being updated by calculating the difference between the number of paths to each descendant in the pair and aggregating the value across term pairs. In case of human proteins, ∼ 38.5% of the values are discarded as we allow only term pairs with non-zero values. The pseudocode in Algorithm 4 demonstrates the computation of SIC scores for all GO term pairs.

##### Algorithm 3 GOpairSICcomp

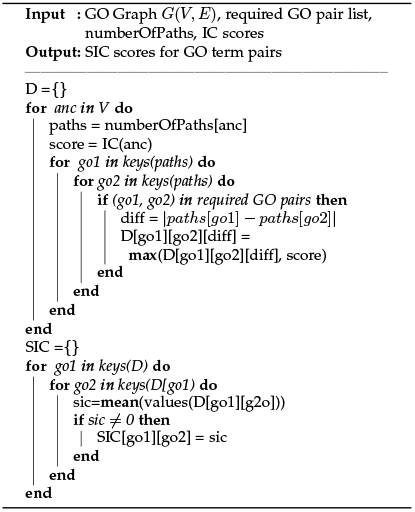

We can now explain the computation of semantic similarity for GO term pairs. For each of the cluster centers *c* of *SG*, the smallest distance to all other GO terms *t* is being computed using Breadth-First Search on *SG* and on its transposition and the minimum of both values is used as *d*(*c, t*) — the smallest distance between *c* and *t*. Then, for all the GO term pairs for which we calculated the *SIC* in the previous point, we compute the *MDMS* and multiply *SIC* and *MDMS* to obtain semantic similarity *sim*. Note that if *SIC* score for a term pair is 0, it is not present in the *SIC* structure, so we don’t compute *sim* for this pair. Additionally, as with *SIC*, we exclude any *sim* scores equal to 0. In case of human proteins, ∼ 0.3% of values are discarded this way. As *SIC, MDMS* and *sim* functions are symmetric, and we store their values for ordered pairs only.

#### 3.5.4 Computation of the semantic similarity for protein pairs

Finally, we are ready to explain the computation of the similarity for each protein pair (see Algorithm 5) which follows the description in Subsection 3.3. First, for each protein pair and all its combinations of GO terms, similarity is being computed using the broadcasted semantic similarity from the previous step. Then, the maximal similarity is found using map-reduce implemented with Spark transformations and used to normalize the results.

#### 3.5.5 Optimizations

In this section we describe optimizations applied in our implementation that had significantly influenced the performance of the computation.

##### Algorithm 4 ProteinPairSimComp

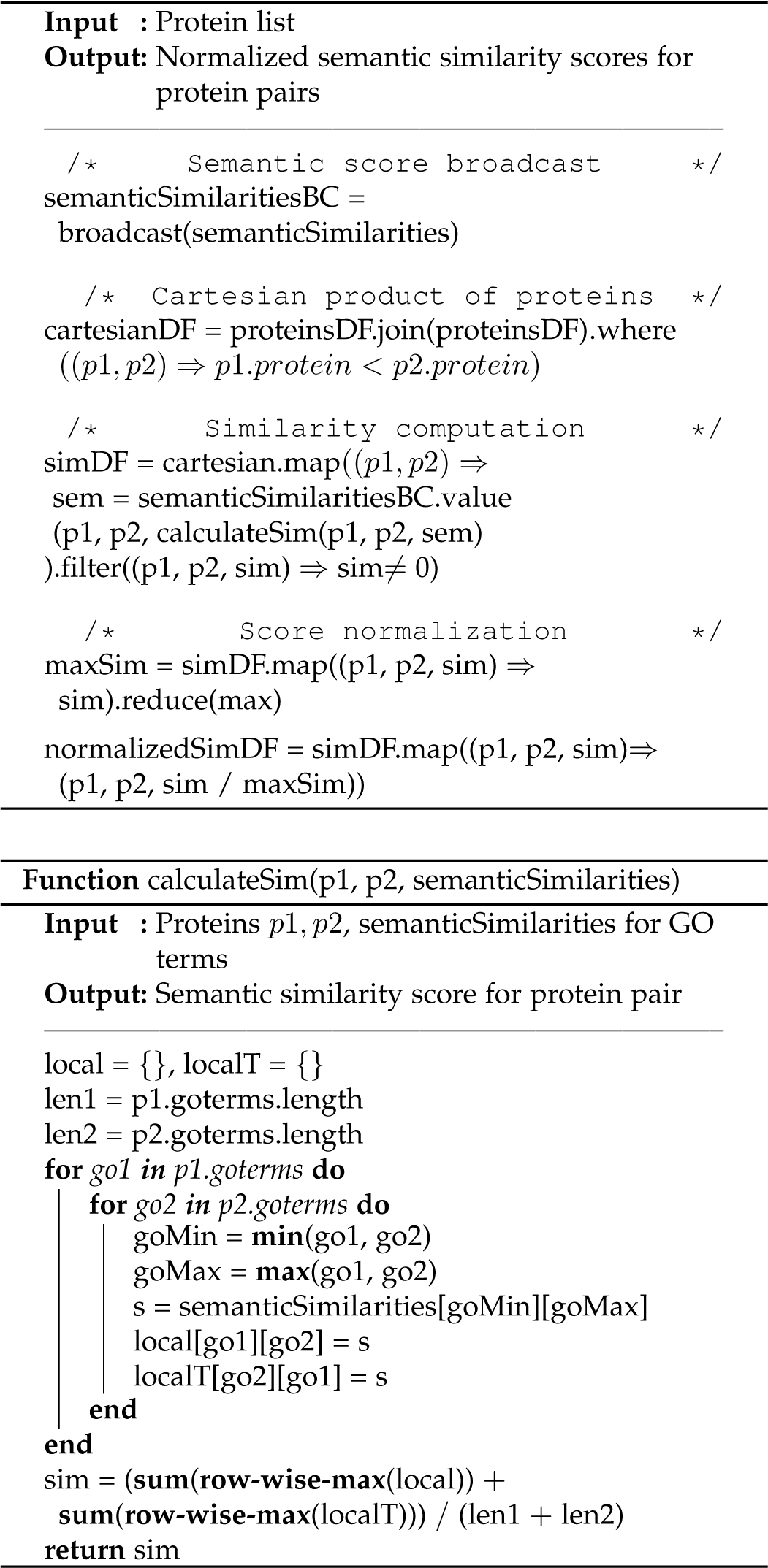

##### Multiple variants at once

In order to compute the fuzzy semantic score for protein pair (*p, q*), similarities 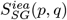 across all GO term subgraphs and annotations need to be combined. Instead of running the algorithm 6 times — once for each subgraph and annotation variant — and going through all its stages, we calculate everything at once and keep track of the variants.

##### Data format

Throughout the whole computation, we encode all the GO terms and variant identifiers as integers instead of strings. This not only decreases the amount of memory needed, but also the footprint of the intra-cluster communication, which saves time.

##### Spark optimizations

The program has been implemented using Spark’s Dataset API, rather than lower-level RDD API. Dataset API is less expressive, as all computations have to be translatable to SQL-like operations on structured data. Yet, this enables several significant optimizations within Spark internal framework [63], [64], such as:

- *Memory Management and Binary Processing:* leveraging application semantics to manage memory explicitly and eliminate the overhead of JVM object model and garbage collection.
- *Cache-aware computation:* Algorithms and data structures to exploit memory hierarchy.
- *Code generation:* Using code generation to exploit modern compilers and CPUs.

## 4 Results and Discussion

We have used complete human proteome datasets and computed fuzzy semantic score of binding affinity for all possible protein pairs. The proposed method is able to quantify the binding affinity of any two proteins within a range of [0,1]. To interpret the numerical result, a threshold cut-off *μ* is used. If the affinity scores is greater than *μ*, it is considered as positive otherwise negative. Comparison of different values of *μ* is shown in Table 3 where for a given value of *μ* we can see the numbers of positive and negative interactions as well as how well the classifier matches the gold sample and whole dataset. The performance evaluation is highlighted in terms of low false positive rate (FPR) and false negative rate (FNR). We also compute: *precision* (a.k.a positive predictive value) which is the fraction of true positives to the total of true and false positives; *recall* (a.k.a sensitivity) which is the fraction of true positives to the total of true positives and false negatives; and area under the curve (AUC) of receiver operating characteristic, which is obtained by plotting the true positive rate against the false positive rate.

**TABLE 3.** Comparison of effects of different values of the threshold cut-off

| $\mu$ | FuzzyPPI | | Gold-PPI (Positive and Negative) | | | | | All-PPI (Positive and Negative) | | | | |
| --- | --- | --- | --- | --- | --- | --- | --- | --- | --- | --- | --- | --- |
|  | Positive | Negative | FPR | FNR | Precision | Recall | AUC | FPR | FNR | Precision | Recall | AUC |
| $10^{-5}$ | 155101018 | 25275503 | 0.963 | 0.009 | 0.673 | 0.991 | 0.514 | 0.763 | 0.041 | 0.889 | 0.959 | 0.598 |
| $10^{-4}$ | 154560195 | 25816326 | 0.962 | 0.01 | 0.674 | 0.99 | 0.514 | 0.762 | 0.042 | 0.889 | 0.958 | 0.598 |
| $10^{-3}$ | 148640177 | 31736344 | 0.95 | 0.012 | 0.676 | 0.988 | 0.519 | 0.752 | 0.053 | 0.889 | 0.947 | 0.598 |
| 0.01 | 120473678 | 59902843 | 0.857 | 0.025 | 0.695 | 0.975 | 0.559 | 0.674 | 0.118 | 0.893 | 0.882 | 0.604 |
| 0.02 | 100445238 | 79931283 | 0.766 | 0.036 | 0.716 | 0.964 | 0.599 | 0.601 | 0.176 | 0.897 | 0.824 | 0.612 |
| 0.03 | 84384625 | 95991896 | 0.673 | 0.05 | 0.739 | 0.95 | 0.638 | 0.528 | 0.232 | 0.903 | 0.768 | 0.62 |
| 0.04 | 70915010 | 109461511 | 0.585 | 0.069 | 0.761 | 0.931 | 0.673 | 0.456 | 0.288 | 0.909 | 0.712 | 0.628 |
| 0.05 | 59805296 | 120571225 | 0.502 | 0.09 | 0.784 | 0.91 | 0.704 | 0.39 | 0.341 | 0.915 | 0.659 | 0.634 |
| 0.06 | 50725562 | 129650959 | 0.429 | 0.113 | 0.806 | 0.887 | 0.729 | 0.332 | 0.392 | 0.921 | 0.608 | 0.636 |
| 0.07 | 43432944 | 136943577 | 0.364 | 0.139 | 0.826 | 0.861 | 0.749 | 0.281 | 0.44 | 0.927 | 0.56 | 0.637 |
| 0.08 | 37209668 | 143166853 | 0.308 | 0.165 | 0.844 | 0.835 | 0.763 | 0.237 | 0.484 | 0.933 | 0.516 | 0.637 |
| 0.09 | 32219592 | 148156929 | 0.261 | 0.191 | 0.861 | 0.809 | 0.774 | 0.2 | 0.526 | 0.938 | 0.474 | 0.638 |
| 0.1 | 27810210 | 152566311 | 0.222 | 0.218 | 0.876 | 0.782 | 0.78 | 0.168 | 0.565 | 0.943 | 0.435 | 0.64 |
| 0.2 | 6774257 | 173602264 | 0.044 | 0.511 | 0.957 | 0.489 | 0.723 | 0.032 | 0.82 | 0.973 | 0.18 | 0.574 |
| 0.3 | 2236133 | 178140388 | 0.01 | 0.718 | 0.983 | 0.282 | 0.636 | 0.007 | 0.92 | 0.987 | 0.08 | 0.537 |
| 0.4 | 872529 | 179503992 | 0.002 | 0.837 | 0.993 | 0.163 | 0.58 | 0.002 | 0.963 | 0.993 | 0.037 | 0.518 |
| 0.5 | 429050 | 179947471 | 0.001 | 0.905 | 0.996 | 0.095 | 0.547 | 0.0008 | 0.982 | 0.996 | 0.018 | 0.509 |
| 0.6 | 182139 | 180194382 | 0 | 0.943 | 0.998 | 0.057 | 0.528 | 0 | 0.991 | 0.998 | 0.009 | 0.505 |
| 0.7 | 80483 | 180296038 | 0 | 0.967 | 0.999 | 0.033 | 0.517 | 0 | 0.995 | 0.999 | 0.005 | 0.503 |
| 0.8 | 24458 | 180352063 | 0 | 0.982 | 1 | 0.018 | 0.509 | 0 | 0.997 | 1 | 0.003 | 0.501 |
| 0.9 | 10184 | 180366337 | 0 | 0.993 | 1 | 0.007 | 0.503 | 0 | 0.999 | 1 | 0.001 | 0.501 |
| 0.95 | 2654 | 180373867 | 0 | 0.997 | 1 | 0.003 | 0.501 | 0 | 1 | 1 | 0 | 0.5 |
| 1 | 0 | 180376521 | 0 | 1 | 1 | 0 | 0.5 | 0 | 1 | 1 | 0 | 0.5 |

### 4.1 Positive High Quality Interaction (PHQI)

We have identified Positive High Quality Interactions (PHQI) with low FPR for *μ* ≥ 0.6. The low FPR score (*type-1* error) indicates that the interactions determined to be positive with *μ* ≥ 0.6 are very reliable, *i.e*., have very-low possibility of being negative. Thus the interactions with increasing threshold from **0.6** onward identifies Positive High Quality Interaction (PHQIs). The significant performance scores are highlighted (green) in the Table 3.

### 4.2 Negative High Quality Interaction (NHQI)

We have also identified Negative High Quality Interaction (NHQI) with low FNR for *μ* ≤10^−3^. The low FNR score (type-2 error) that the interactions determined to be negative with *μ* = ≤10^−3^ are very reliable, *i.e*., have very low probability of being positive. The proposed approach has achieved low FNR (< 0.01) on Gold-PPI dataset at *μ* = 10^−3^ threshold. However, in All-PPI dataset the FNR score is 0.053 at the mentioned threshold. The thresholds with significant FNR scores are highlighted (orange) in the Table 3.

### 4.3 Critical threshold

The experimental results from Table 3, suggest that the PHQI and NHQI interactions can be successfully identified using proposed scoring scheme (*μ* ≤10^−3^ for NHQI and *μ* ≥0.6 for PHQI). With the increasing values of threshold the quality of negative interactions gradually decreases. Similarly, the quality of the positive interactions drops as the threshold is decreased. To impose a critical cutoff on the scoring, different evaluation metrics (precision, recall and AUC) have been used. The precision scores show improving trend with the increase of threshold whereas recall score shows a better performance at lower threshold. AUC score became crucial to fix the threshold point, where both the precision and recall are on balance. Based on the AUC scores on both datasets (ALL-PPI and Gold-PPI), the 0.1 is considered as the critical threshold. With the threshold cut-off of 0.1, for both datasets we have the highest AUC value (0.64 in All-PPI and 0.78 in Gold-PPI).

### 4.4 Validation with respect to *state-of-the-art*

The proposed approach is able to provide a fuzzy semantic score between any protein pair that signifies interaction affinity between them. The biological significance of these scoring scheme is to categorise the interaction space into different significant levels such as high confidence (both positive and negative interaction) and low confidence (uncertain).

This categorisation has significant effect on machine learning based PPI prediction models. To establish the importance of the proposed scheme, three different datasets are selected from the interaction data pool using above described scoring cutoffs and PPI prediction performances have been evaluated using two independent PPI prediction methods, *JUPPI* [65], and *Ding et al*. [66]. First, ***PHQI-NHQI*** dataset, is selected from high quality positive (with *μ* ≥0.6) and high quality negative (with *μ* ≤10^−3^). Second, ***AP-AN*** dataset, is selected randomly from the pool of all known positive and negative interaction data. Finally, a ***UP-UN*** dataset is selected from the uncertain region of interaction space (*μ* < 0.6 and *μ* > 10^−3^) as shown in Table 3. In this final dataset, all ambiguous interactions are removed from both positive and negative interaction sets for clarity of the test. The performances of both methods are evaluated with five statistical metrics (AUC, AUPRC, ACCU, F1 and MCC) and scores are reported in Table 4. Both methods have shown a significant performance improvement on ***PHQI-NHQI*** dataset compared to ***AP-AN*** and ***UP-UN***. The performances on ***UP-UN*** dataset, is worst than other two due to the uncertain positive and negative selection in training data. The AUC and AUPRC curves from three sets of interaction dataset are shown in Figure 4.

**TABLE 4.** Performance analysis of **JUPPI** and **Ding *et al***. with three different dataset using Fuzzy semantic scoring threshold.

| Method | Data Type | AUC | AUPRC | ACC | F1 | MCC |
| --- | --- | --- | --- | --- | --- | --- |
| JUPPI [65] | PHQI-NHQI | 0.996 | 0.996 | 0.973 | 0.973 | 0.946 |
|  | AP-AN | 0.972 | 0.974 | 0.910 | 0.908 | 0.823 |
|  | UP-UN | 0.939 | 0.940 | 0.868 | 0.865 | 0.738 |
| Ding *et al.* [66] | PHQI-NHQI | 0.924 | 0.927 | 0.836 | 0.832 | 0.672 |
|  | AP-AN | 0.856 | 0.857 | 0.767 | 0.763 | 0.535 |
|  | UP-UN | 0.836 | 0.826 | 0.747 | 0.737 | 0.496 |

**Fig. 4.**
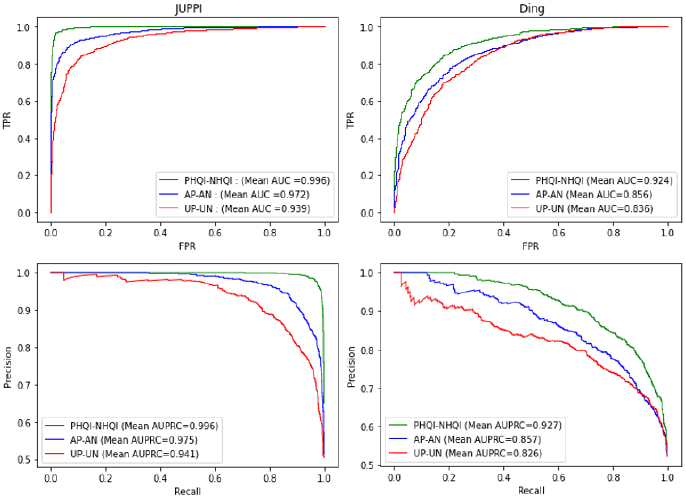
Performance changes with respect to AUC and AUPRC on three set of PPI data extracted from PHQI-NHQI, AAP-AN and UP-UN. Plots in the 1^*st*^ row represents AUC, and the 2^*nd*^ row represent AUPRC, respectively. The 1^*st*^ and 2^*nd*^, column-wise plots show the curves from JUPPI [65] and Ding *et al*. [66].

### 4.5 Significance of FSS at proteome level disease analysis

In this section we have extracted significance sub clusters from complete fuzzy semantic network of human proteome. For graph sub cluster retrieval, MCL clustering algorithm [67], [69] has been employed on the complete human interactome with developed fuzzy semantic scores. To evaluate the significance of the developed Fuzzy PPI model, we characterise these sub clusters with respect to the disease association with twenty groups of human cancer disease from the Pathology Atlas of Human Protein Atlas [68]. The cluster-disease association is presented with an example cluster on three cutt-off thresholds (*μ*) of FSS (≥ 0.5, ≥ 0.7 and ≥0.9). The cluster hierarchy and their association is shown in Figure 5.

**Fig. 5.**
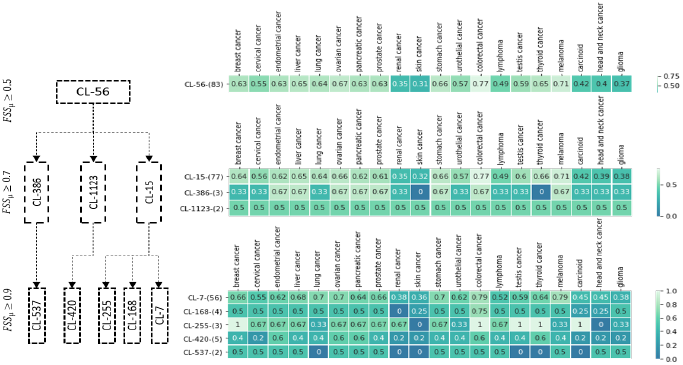
Cluster hierarchy and heatmap representation of clusters extracted at thee *μ* thresholds (≥ 0.5, ≥ 0.7 and ≥ 0.9) of FSS using MCL algorithm [67]. The significance of the clusters are highlighted with different cancer diseases association. Scores within the heatmap cell represents the percentage of protein association with respect to cancer disease type. Protein to cancer association (high confidence) are retrieved from Human Protein Atlas [68].

## 5 Conclusion

In this paper we have present a new fuzzy affinity based scoring scheme for prediction of interaction affinity pairs of human proteins. The work is built upon the GO associations of respective proteins and the underlying GO graphs for MF, CC and BP subgraphs. A graph clustering approach has been used to identify representative cluster centers in a graph, and the ancestor-descendent relationships between two nodes has been utilized to design the fuzzy affinity functions. One of the major limitations of the work is that it depends heavily on the GO annotations of the respective proteins. In cases, where GO annotations are not available, the interaction affinity cannot be predicted. However, from among 20 350 reviewed human proteins, only 1 356 proteins did not have a matching GO annotation, and had to be excluded from our study, which allowed us to successfully estimated interaction affinity between ∼ 180 million of protein pairs. Also, the GO information/annotation and underlying GO graphs get updated periodically. Minor perturbation in the GO network may lead to changes in the PPI affinity scores. Therefore, we also propose to update the web-server periodically with new releases of the GO annotations. Computation of such a large number of interaction estimations would not have been possible without use of a Spark based parallelization. As for comparison, the work by Dutta *et al*. [51] used only 4 726 interactions from ∼ 2000 human proteins. With a standard desktop computer and sequential implementation, the graph clustering and affinity assessment score estimation algorithms take days/weeks to complete. In contrast, our ∼ 180 million interaction estimations took less than a week in a Spark based parallel cluster setup. We plan to extend our work with both reviewed and unreviewed proteins and also for multi-organism PPI prediction problems, leading to more than trillion interactions. Finally, one of the major objectives of our work was to qualitatively categorize the protein interactions, based on the affinity scores. It may be observed that the Fuzzy-PPI may be useful for identifying high quality positive interactions with very low FPR, as well as high quality negative data selection with very low FPR. Our work has also been compared with the *state-of-the-art* in the domain and the its effectiveness has been validated.

## Acknowledgments

This work is partially supported by the CMATER research laboratory of the Computer Science and Engineering Department, Jadavpur University, India, PURSE-II and UPE-II grants. **SB**:Department of Biotechnology grant (BT/PR16356/BID/7/596/2016), Government of India.

---

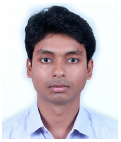

**Anup Kumar Halder** (S’17) is an Assistant Professor in the Department of Computer Science & Engineering, University of Engineering & Management, Kolkata 700156, West Bengal, India. He received his B.Tech and M.E. degree in Computer Science and Engineering from Bankura Unnayani Institute of Engineering in 2011 and Jadavpur University in 2014 respectively. He received his Ph.D. in Engineering degree from Jadavpur University in 2021. He is the recipient of Visvesvaraya PhD fellowship from MeitY, Govt. of India (2015-2020). His area of current research interest includes computational biology and bioinformatics, genomic and proteomic network analysis, pattern recognition and machine learning.

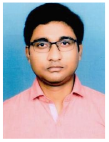

**Soumyendu Sekhar Bandyopadhyay** is an Assistant Professor in the Department of Computer Science & Engineering, School of Engineering and Technology, Adamas University, Kolkata 700126, West Bengal, India. He is working towards his PhD degree in the Department of Computer Science and Engineering, Jadavpur University, Kolkata, India. He received his MCA degree from Bengal Institute of Technology, Kolkata, India in 2009 and M.Tech degree from University of Calcutta, Kolkata, India in 2012. His current research interest is bioinformatics and large scale biological networks.

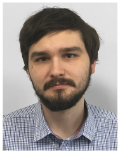

**Witold Jedrzejewski** is working towards his PhD degree at the Faculty of Mathematics, Informatics and Mechanics, University of Warsaw, Poland. He received his MSc in informatics from the Institute of Informatics, University of Warsaw in 2018. His current research interest includes Massively Parallel Computation and algorithms for distributed data processing. He also works as a consultant for ad-tech and computer networking industries in matters of machine learning and big data engineering.

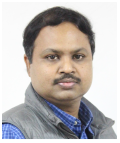

**Subhadip Basu** (SM’12) is a Professor in the Department of Computer Science and Engineering, Jadavpur University, India. He received PhD in Engineering from Jadavpur University in 2006 and has more than 200 research publications in notable Journals and Conferences in the areas of Bioinformatics, Image Analysis, Pattern Recognition etc. Dr. Basu is the recipient of DAAD fellowship from Germany, BOYSCAST fellowship and UGC Research Award from the Govt. of India, HIVIP fellowship from Hitachi, Japan, EMMA and cLink fellowships from the European Union.

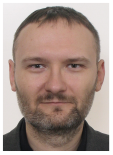

**Jacek Sroka** Jacek Sroka is an assistant professor at Faculty of Mathematics, Informatics and Mechanics, University of Warsaw, Poland. He got his PhD in informatics in 2009 from the University of Warsaw. His research interests are: data bases, cluster computing, Big data, bioinformatics and workflow modelling. He is also a consultant for the National Electora Office of Republic of Poland and helps to organize the national elections.

## References

[1] U. Stelzl, U. Worm, M. Lalowski, C. Haenig, F. H. Brembeck, H. Goehler, M. Stroedicke, M. Zenkner, A. Schoenherr, S. Koeppen et al., “A human protein-protein interaction network: a resource for annotating the proteome,” Cell, vol. 122, no. 6, pp. 957–968, 2005.

[2] T. Nepusz, H. Yu, and A. Paccanaro, “Detecting overlapping protein complexes in protein-protein interaction networks,” Nature methods, vol. 9, no. 5, p. 471, 2012.

[3] A. Vazquez, A. Flammini, A. Maritan, and A. Vespignani, “Global protein function prediction from protein-protein interaction networks,” Nature biotechnology, vol. 21, no. 6, pp. 697–700, 2003.

[4] Y.-R. Cho, Y. Xin, and G. Speegle, “P-finder: Reconstruction of signaling networks from protein-protein interactions and go annotations,” IEEE/ACM transactions on computational biology and bioinformatics, vol. 12, no. 2, pp. 309–321, 2014.

[5] G. Bebek and J. Yang, “Pathfinder: mining signal transduction pathway segments from protein-protein interaction networks,” BMC bioinformatics, vol. 8, no. 1, p. 335, 2007.

[6] A. K. Halder, M. Denkiewicz, K. Sengupta, S. Basu, and D. Plewczynski, “Aggregated network centrality shows nonrandom structure of genomic and proteomic networks,” Methods, 2019.

[7] A. K. Halder, P. Dutta, M. Kundu, S. Basu, and M. Nasipuri, “Review of computational methods for virus–host protein interaction prediction: a case study on novel ebola–human interactions,” Briefings in functional genomics, vol. 17, no. 6, pp. 381–391, 2018.

[8] M. Ay, K.-I. Goh, M. E. Cusick, A.-L. Barabasi, M. Vidal et al., “Drug–target network,” Nature biotechnology, vol. 25, no. 10, pp. 1119–1127, 2007.

[9] H. Ruffner, A. Bauer, and T. Bouwmeester, “Human protein– protein interaction networks and the value for drug discovery,” Drug discovery today, vol. 12, no. 17-18, pp. 709–716, 2007.

[10] D. Dasagrandhi, A. S. K. Ravindran, A. Muthuswamy, and K. Jayachandran, “Construction and analysis of protein-protein interaction network: Role in identification of key signaling molecules involved in a disease pathway,” in Computer Applications in Drug Discovery and Development. IGI Global, 2019, pp. 204–220.

[11] Y.-C. Chen, S. V. Rajagopala, T. Stellberger, and P. Uetz, “Exhaustive benchmarking of the yeast two-hybrid system,” Nature methods, vol. 7, no. 9, pp. 667–668, 2010.

[12] S. Fields, “High-throughput two-hybrid analysis: The promise and the peril,” The FEBS journal, vol. 272, no. 21, pp. 5391–5399, 2005.

[13] A.-C. Gavin, M. Bösche, R. Krause, P. Grandi, M. Marzioch, A. Bauer, J. Schultz, J. M. Rick, A.-M. Michon, C.-M. Cruciat et al., “Functional organization of the yeast proteome by systematic analysis of protein complexes,” Nature, vol. 415, no. 6868, pp. 141–147, 2002.

[14] Y. Ho, A. Gruhler, A. Heilbut, G. D. Bader, L. Moore, S.-L. Adams, A. Millar, P. Taylor, K. Bennett, K. Boutilier et al., “Systematic identification of protein complexes in saccharomyces cerevisiae by mass spectrometry,” Nature, vol. 415, no. 6868, pp. 180–183, 2002.

[15] M. Bellucci, F. Agostini, M. Masin, and G. G. Tartaglia, “Predicting protein associations with long noncoding rnas,” Nature methods, vol. 8, no. 6, p. 444, 2011.

[16] Y. Guo, L. Yu, Z. Wen, and M. Li, “Using support vector machine combined with auto covariance to predict protein–protein interactions from protein sequences,” Nucleic acids research, vol. 36, no. 9, pp. 3025–3030, 2008.

[17] T. Hamp and B. Rost, “Evolutionary profiles improve protein– protein interaction prediction from sequence,” Bioinformatics, vol. 31, no. 12, pp. 1945–1950, 2015.

[18] V. Perovic, N. Sumonja, L. A. Marsh, S. Radovanovic, M. Vukicevic, S. G. Roberts, and N. Veljkovic, “Idppi: Protein-protein interaction analyses of human intrinsically disordered proteins,” Scientific reports, vol. 8, no. 1, pp. 1–10, 2018.

[19] P. Chatterjee, S. Basu, M. Kundu, M. Nasipuri, and D. Plewczynski, “Ppi svm: Prediction of protein-protein interactions using machine learning, domain-domain affinities and frequency tables,” Cellular and Molecular Biology Letters, vol. 16, no. 2, pp. 264–278, 2011.

[20] K. K. Wan, J. Park, and J. K. Suh, “Large scale statistical prediction of protein-protein interaction by potentially interacting domain (pid) pair,” Genome Informatics, vol. 13, pp. 42–50, 2002.

[21] S. Bandyopadhyay and K. Mallick, “A new feature vector based on gene ontology terms for protein-protein interaction prediction,” IEEE/ACM transactions on computational biology and bioinformatics, vol. 14, no. 4, pp. 762–770, 2016.

[22] S. R. Maetschke, M. Simonsen, M. J. Davis, and M. A. Ragan, “Gene ontology-driven inference of protein–protein interactions using inducers,” Bioinformatics, vol. 28, no. 1, pp. 69–75, 2012.

[23] A. Ben-Hur and W. S. Noble, “Kernel methods for predicting protein–protein interactions,” Bioinformatics, vol. 21, no. suppl 1, pp. i38–i46, 2005.

[24] P. Blohm, G. Frishman, P. Smialowski, F. Goebels, B. Wachinger, A. Ruepp, and D. Frishman, “Negatome 2.0: a database of noninteracting proteins derived by literature mining, manual annotation and protein structure analysis,” Nucleic acids research, vol. 42, no. D1, pp. D396–D400, 2014.

[25] L. G. Trabuco, M. J. Betts, and R. B. Russell, “Negative protein– protein interaction datasets derived from large-scale two-hybrid experiments,” Methods, vol. 58, no. 4, pp. 343–348, 2012.

[26] U. Consortium, “Uniprot: a hub for protein information,” Nucleic acids research, vol. 43, no. D1, pp. D204–D212, 2015.

[27] J. H. Moon, S. Lim, K. Jo, S. Lee, S. Seo, and S. Kim, “Pintnet: construction of condition-specific pathway interaction network by computing shortest paths on weighted ppi,” BMC systems biology, vol. 11, no. 2, p. 15, 2017.

[28] C. Zhao and Z. Wang, “Gogo: an improved algorithm to measure the semantic similarity between gene ontology terms,” Scientific reports, vol. 8, no. 1, pp. 1–10, 2018.

[29] G. O. Consortium, “The gene ontology resource: 20 years and still going strong,” Nucleic acids research, vol. 47, no. D1, pp. D330–D338, 2019.

[30] I. Kuznetsova, A. Lugmayr, S. J. Siira, O. Rackham, and A. Filipovska, “Cirgo: an alternative circular way of visualising gene ontology terms,” BMC bioinformatics, vol. 20, no. 1, pp. 1–7, 2019.

[31] J. Zhu, Q. Zhao, E. Katsevich, and C. Sabatti, “Exploratory gene ontology analysis with interactive visualization,” Scientific reports, vol. 9, no. 1, pp. 1–9, 2019.

[32] H. Hassan and S. Shanak, “Gotrapper: a tool to navigate through branches of gene ontology hierarchy,” BMC bioinformatics, vol. 20, no. 1, pp. 1–6, 2019.

[33] M. Armbrust, R. S. Xin, C. Lian, Y. Huai, D. Liu, J. K. Bradley, X. Meng, T. Kaftan, M. J. Franklin, A. Ghodsi, and M. Zaharia, “Spark SQL: relational data processing in spark,” in Proceedings of the 2015 ACM SIGMOD International Conference on Management of Data, Melbourne, Victoria, Australia, May 31 - June 4, 2015, T. K. Sellis, S. B. Davidson, and Z. G. Ives, Eds. ACM, 2015, pp. 1383–1394. [Online]. Available: https://doi.org/10.1145/2723372.2742797

[34] M. Zaharia, M. Chowdhury, T. Das, A. Dave, J. Ma, M. McCauly, M. J. Franklin, S. Shenker, and I. Stoica, “Resilient distributed datasets: A fault-tolerant abstraction for in-memory cluster computing,” in Proceedings of the 9th USENIX Symposium on Networked Systems Design and Implementation, NSDI 2012, San Jose, CA, USA, April 25-27, 2012, S. D. Gribble and D. Katabi, Eds. USENIX Association, 2012, pp. 15–28. [Online]. Available: https://www.usenix.org/conference/nsdi12/technical-sessions/presentation/zaharia

[35] V. Pekar and S. Staab, “Taxonomy learning-factoring the structure of a taxonomy into a semantic classification decision,” in COLING 2002: The 19th International Conference on Computational Linguistics, 2002.

[36] Z. Wu and M. Palmer, “Verbs semantics and lexical selection,” in Proceedings of the 32nd annual meeting on Association for Computational Linguistics. Association for Computational Linguistics, 1994, pp. 133–138.

[37] H. Yu, L. Gao, K. Tu, and Z. Guo, “Broadly predicting specific gene functions with expression similarity and taxonomy similarity,” Gene, vol. 352, pp. 75–81, 2005.

[38] C. Pesquita, D. Faria, A. O. Falcao, P. Lord, and F. M. Couto, “Semantic similarity in biomedical ontologies,” PLoS computational biology, vol. 5, no. 7, 2009.

[39] M. G. Ahsaee, M. Naghibzadeh, and S. E. Y. Naeini, “Semantic similarity assessment of words using weighted wordnet,” International Journal of Machine Learning and Cybernetics, vol. 5, no. 3, pp. 479–490, 2014.

[40] P. Resnik, “Semantic similarity in a taxonomy: An information-based measure and its application to problems of ambiguity in natural language,” Journal of artificial intelligence research, vol. 11, pp. 95–130, 1999.

[41] D. Lin et al., “An information-theoretic definition of similarity.” In Icml, vol. 98, no. 1998, 1998, pp. 296–304.

[42] J. J. Jiang and D. W. Conrath, “Semantic similarity based on corpus statistics and lexical taxonomy,” arXiv preprint cmp-lg/9709008, 1997.

[43] A. Schlicker, F. S. Domingues, J. Rahnenführer, and T. Lengauer, “A new measure for functional similarity of gene products based on gene ontology,” BMC bioinformatics, vol. 7, no. 1, p. 302, 2006.

[44] C. E. Shannon, “A mathematical theory of communication,” Bell system technical journal, vol. 27, no. 3, pp. 379–423, 1948.

[45] X. Wu, E. Pang, K. Lin, and Z.-M. Pei, “Improving the measurement of semantic similarity between gene ontology terms and gene products: insights from an edge-and ic-based hybrid method,” PloS one, vol. 8, no. 5, 2013.

[46] G. K. Mazandu and N. J. Mulder, “Information content-based gene ontology semantic similarity approaches: toward a unified framework theory,” BioMed research international, vol. 2013, 2013.

[47] F. M. Couto and M. J. Silva, “Disjunctive shared information between ontology concepts: application to gene ontology,” Journal of biomedical semantics, vol. 2, no. 1, p. 5, 2011.

[48] P. H. Guzzi, M. Mina, C. Guerra, and M. Cannataro, “Semantic similarity analysis of protein data: assessment with biological features and issues,” Briefings in bioinformatics, vol. 13, no. 5, pp. 569–585, 2012.

[49] G. K. Mazandu, E. R. Chimusa, M. Mbiyavanga, and N. J. Mulder, “A-dago-fun: an adaptable gene ontology semantic similarity-based functional analysis tool,” Bioinformatics, vol. 32, no. 3, pp. 477–479, 2016.

[50] A. Nagar and H. Al-Mubaid, “A hybrid semantic similarity measure for gene ontology based on offspring and path length,” in 2015 IEEE Conference on Computational Intelligence in Bioinformatics and Computational Biology (CIBCB). IEEE, 2015, pp. 1–7.

[51] P. Dutta, S. Basu, and M. Kundu, “Assessment of semantic similarity between proteins using information content and topological properties of the gene ontology graph,” IEEE/ACM transactions on computational biology and bioinformatics, vol. 15, no. 3, pp. 839–849, 2017.

[52] A. Bairoch and R. Apweiler, “The swiss-prot protein sequence database and its supplement trembl in 2000,” Nucleic acids research, vol. 28, no. 1, pp. 45–48, 2000.

[53] C. O’Donovan, M. J. Martin, A. Gattiker, E. Gasteiger, A. Bairoch, and R. Apweiler, “High-quality protein knowledge resource: Swiss-prot and trembl,” Briefings in bioinformatics, vol. 3, no. 3, pp. 275–284, 2002.

[54] C. H. Wu, L.-S. L. Yeh, H. Huang, L. Arminski, J. Castro-Alvear, Y. Chen, Z. Hu, P. Kourtesis, R. S. Ledley, B. E. Suzek et al., “The protein information resource,” Nucleic acids research, vol. 31, no. 1, pp. 345–347, 2003.

[55] S. Pundir, M. Magrane, M. J. Martin, C. O’Donovan, and U. Consortium, “Searching and navigating uniprot databases,” Current protocols in bioinformatics, vol. 50, no. 1, pp. 1–27, 2015.

[56] M. H. Schaefer, J.-F. Fontaine, A. Vinayagam, P. Porras, E. E. Wanker, and M. A. Andrade-Navarro, “Hippie: Integrating protein interaction networks with experiment based quality scores,” PloS one, vol. 7, no. 2, 2012.

[57] D. Szklarczyk, A. L. Gable, D. Lyon, A. Junge, S. Wyder, J. Huerta-Cepas, M. Simonovic, N. T. Doncheva, J. H. Morris, P. Bork et al., “String v11: protein–protein association networks with increased coverage, supporting functional discovery in genome-wide experimental datasets,” Nucleic acids research, vol. 47, no. D1, pp. D607–D613, 2019.

[58] R. Oughtred, C. Stark, B.-J. Breitkreutz, J. Rust, L. Boucher, C. Chang, N. Kolas, L. O’Donnell, G. Leung, R. McAdam et al., “The biogrid interaction database: 2019 update,” Nucleic acids research, vol. 47, no. D1, pp. D529–D541, 2019.

[59] I. Xenarios, D. W. Rice, L. Salwinski, M. K. Baron, E. M. Marcotte, and D. Eisenberg, “Dip: the database of interacting proteins,” Nucleic acids research, vol. 28, no. 1, pp. 289–291, 2000.

[60] K. Luck, D.-K. Kim, L. Lambourne, K. Spirohn, B. E. Begg, W. Bian, R. Brignall, T. Cafarelli, F. J. Campos-Laborie, B. Charloteaux et al., “A reference map of the human binary protein interactome,” Nature, pp. 1–7, 2020.

[61] H.-S. Park and C.-H. Jun, “A simple and fast algorithm for k-medoids clustering,” Expert Systems with Applications, vol. 36, no. 2, Part 2, pp. 3336–3341, 2009. [Online]. Available: https://www.sciencedirect.com/science/article/pii/S095741740800081X

[62] J. Beel, B. Gipp, S. Langer, and C. Breitinger, “Research-paper recommender systems: a literature survey,” International Journal on Digital Libraries, vol. 17, no. 4, pp. 305–338, 2016. [Online]. Available: http://dx.doi.org/10.1007/s00799-015-0156-0

[63] M. Armbrust, T. Das, A. Davidson, A. Ghodsi, A. Or, J. Rosen, I. Stoica, P. Wendell, R. Xin, and M. Zaharia, “Scaling spark in the real world: Performance and usability,” Proc. VLDB Endow., vol. 8, no. 12, p. 1840–1843, Aug. 2015. [Online]. Available: https://doi.org/10.14778/2824032.2824080

[64] J. Rosen and R. Xin, “Packaging - google chrome,” https://databricks.com/blog/2015/04/28/project-tungsten-bringing-spark-closer-to-bare-metal.html.

[65] A. K. Halder, S. S. Bandyopadhyay, P. Chatterjee, M. Nasipuri, D. Plewczynski, and S. Basu, “Juppi: A multi-level feature based method for ppi prediction and a refined strategy for performance assessment,” IEEE/ACM Transactions on Computational Biology and Bioinformatics, 2020.

[66] Y. Ding, J. Tang, and F. Guo, “Predicting protein-protein interactions via multivariate mutual information of protein sequences,” BMC bioinformatics, vol. 17, no. 1, p. 398, 2016.

[67] S. Van Dongen, A new cluster algorithm for graphs. Citeseer, 1998.

[68] M. Uhlen, C. Zhang, S. Lee, E. Sjöstedt, L. Fagerberg, G. Bidkhori, R. Benfeitas, M. Arif, Z. Liu, F. Edfors et al., “A pathology atlas of the human cancer transcriptome,” Science, vol. 357, no. 6352, 2017.

[69] S. Van Dongen, “A stochastic uncoupling process for graphs,” in NATIONAL RESEARCH INSTITUTE FOR MATHEMATICS AND COMPUTER SCIENCE IN THE. Citeseer, 2000.

